# QC-espresso: Brewing quality in proteomics

**DOI:** 10.64898/2026.10.29.762921

**Authors:** Guilherme Pauperio Lanfredi, Jackson Gabriel Miyamoto

**Author notes:** Corresponding author and tool maintainer.

## Abstract

Data-independent acquisition (DIA) proteomics has become a standard approach for high-throughput proteomics, with DIA-NN emerging as one of the most widely adopted data-analysis platforms. However, despite the detailed information available in DIA-NN output reports, proteomics researchers may lack a streamlined and reproducible approach for assessing acquisition quality, identification consistency, and quantitative reliability across experiments. QC-espresso addresses this need through an automated, report-centered quality-control workflow implemented in R and Quarto. The tool processes a DIA-NN report.parquet file together with the corresponding FASTA database and generates a structured HTML report containing plots, run-level statistics, and exportable summary tables. Application to three experimental datasets allowed the visualization of lower identification depth related to high contaminant content; a chromatographic transition accompanied by altered retention-time and peak-width behavior; and the co-occurrence of systematic mass-error bias with changes in precursor- and protein-level quantitative-quality metrics. QC-espresso can be executed directly in R or through a portable Windows distribution with a graphical launcher, with both modes using the same analytical modules and report-generation workflow. The resulting report provides a consistent record for local inspection and troubleshooting of quality-control metrics within individual DIA proteomics experiments. QC-espresso is freely available at https://github.com/lanfredig/QC-espresso_DIA.

## Introduction

The increasing adoption of data-independent acquisition (DIA) mass spectrometry has improved the depth, reproducibility, and quantitative accuracy of proteomics experiments. Among available software solutions, DIA-NN has become a leading platform for DIA data analysis by combining neural-network-based peptide identification, interference correction, and quantitative signal-processing strategies ^1^. DIA-NN also provides confidence and quantitative-quality fields that can support filtering and the evaluation of analytical consistency. Quality-control assessment can support verification after instrument calibration or maintenance, including the examination of replicate acquisitions. As proteomics cohorts scale to hundreds of LC-MS/MS runs, core facilities and research labs also require systematic QC to identify run-to-run drift.

Although DIA-NN provides extensive output tables and quality metrics, these data are distributed across extensive report files and commonly require custom scripting and bioinformatics expertise for systematic interpretation. Many proteomics software tools focus primarily on filtering, normalization, and differential abundance analysis, whereas fewer tools consolidate run performance, sample quality, and acquisition issues into a persistent experiment-level report. Some software environments provide related quality-control capabilities but are distributed under commercial licenses, which may limit their adoption in settings that depend on openly available computational tools.

### From Instrument Monitoring to Experiment-Level Quality Assessment

Existing quality-control tools address important but distinct needs across proteomics workflows. QCloud2 ^2^ provides longitudinal monitoring of LC-MS performance across instruments and laboratories, whereas QSample ^3^ monitors quality indicators from experimental samples while data acquisition is still in progress, supporting timely decisions before an acquisition sequence has been completed. At the postprocessing stage, pmultiqc ^4^ provides scalable and metadata-aware quality-control reporting across multiple proteomics pipelines, including DIA-NN, while QC4DIANN ^5^ provides an interactive Shiny environment for exploring identification and quantitative-quality measurements from DIA-NN results. Together, these tools provide clear advantages for longitudinal instrument surveillance, acquisition-time intervention, multiplatform reporting, and interactive investigation of processed data.

The mode of execution is also relevant to the routine use of quality-control software. Depending on the tool and deployment model, local execution may require programming experience, dependency management, workflow configuration, or previously established computational infrastructure. Although these requirements can be accommodated in centralized facilities and large-scale studies, they may complicate implementation on local data-processing workstations. QC-espresso therefore provides both direct execution in R and a portable Windows distribution with a graphical launcher, allowing the workflow to be used with or without a separately configured R and Quarto environment.

To address these analytical and operational needs, we developed QC-espresso, an R- and Quarto-based workflow that processes DIA-NN report.parquet files together with the corresponding FASTA database. QC-espresso consolidates information related to run performance, sample quality, and acquisition behavior into a persistent HTML report, allowing individual measurements to be interpreted in relation to broader patterns in the data. This report-centered approach complements the longitudinal monitoring, acquisition-time assessment, multipipeline reporting, and interactive exploration provided by existing tools. The use of the corresponding FASTA database further extends the assessment by supporting protein annotation and sequence-coverage analysis.

## Experimental Section

### QC-espresso Architecture and Execution

QC-espresso was implemented in R as a modular workflow comprising a scientific runner, analytical R scripts, and a parameterized Quarto report. The runner manages the execution and passes the validated input and output paths to the Quarto document, which calls the analytical modules in a defined sequence and organizes their graphical and tabular results, while the modular structure allows individual analytical components to be inspected or modified independently.

Two execution modes are available. The R-based mode runs directly from the project environment, whereas the portable Windows mode uses a graphical launcher to collect the input paths and call the same analytical runner. The portable distribution includes the R runtime, package library, and Quarto installation required for execution.

Each execution receives a DIA-NN report.parquet file and its corresponding FASTA database. After input validation and analytical processing, the workflow uses Quarto to render a self-contained HTML report, while run-level Figures of Merit and peptide-supported protein-coverage results are exported as CSV files. The outputs are stored in an independent timestamped execution directory, maintaining their association and preventing previous results from being overwritten.

### Input Validation and Data Processing

QC-espresso first verifies that the selected report.parquet and FASTA files are accessible. The DIA-NN report is then inspected for the fields required by the core workflow and each conditional module. Modules dependent on unavailable or invalid optional measurements are omitted without interrupting generation of the remaining report.

The workflow excludes decoy entries and retains protein-group observations with PG.Q.Value ≤ 0.01. Runs for which no observations remain after filtering are identified in the report and omitted from the corresponding downstream calculations.

Protein-sequence coverage is calculated by matching peptide evidence associated with each DIA-NN Protein.Group to compatible entries in the corresponding FASTA database. Coverage is reported separately for each matched protein sequence without redistributing the quantitative signal assigned by DIA-NN.

Filtered observations are organized according to the analytical level required by each module. Distinct precursor, peptide, and protein-group identifiers are used for identification summaries, whereas quantitative values support signal-composition and intensity analyses. Run identifiers are preserved throughout processing to maintain correspondence between graphical and numerical results.

### Analytical Modules and Quality-Control Outputs

QC-espresso organizes the analysis into modules that receive specific DIA-NN fields and return graphical or numerical outputs to the Quarto report. Modules that depend on optional fields are included only when compatible and valid measurements are available, allowing the remaining report to be generated without assigning values to unsupported measurements.

Among the DIA-NN-derived measurements, Quantity.Quality, Empirical.Quality, and PG.MaxLFQ.Quality are retained without recalculation and presented as complementary views of quantitative reliability. These measurements represent related but non-equivalent aspects of quantitative consistency and are displayed alongside the results from the other analytical modules. Reference lines were included only as visual interpretation aids.

To support run-level review, selected results from these modules are summarized in the Figures of Merit table, providing a concise numerical overview of each run. The measurements remain separate, allowing their individual behavior to be examined in the context of the complete report rather than being reduced to a composite score or automatic quality classification. The inputs, principal operations, and outputs of each module are summarized in Table 1.

**Table 1.** Analytical modules, required inputs, principal operations, and outputs implemented in QC-espresso.

| **Module** | **Input** | **Operations** | **Output** |
|---|---|---|---|
| Identification performance | Precursor-, peptide-, and protein-group identifiers | Run-level counting and comparison across acquisitions | Identification summary tables and plots at precursors, peptide and protein groups levels, including mean, maximum, and total reference values |
| Quantitative signal composition | Protein groups, precursor quantities, and annotations | Aggregation of signal contributions and contaminant fractions | Protein-group composition and contaminant-signal plots, including dominant annotation labels and the total contaminant contribution per run |
| Sequence coverage | Peptide evidence and corresponding FASTA sequences | Peptide-to-sequence mapping and covered-residue calculation | Protein-coverage density plot, Top 100 coverage table, and complete CSV export containing sequence coverage and supporting peptide evidence |
| Peptide-level quality | Charge states, peptide sequences, and cleavage information | Charge state, peptide lengths, and missed-cleavage frequency assessment | Charge-state, peptide-length, and missed-cleavage plots with run-level distributions and dataset-wide reference summaries |
| Chromatographic performance | Retention time, FWHM, peak boundaries, and reference peptides | Peak performance metrics and identifications across gradient assessment | Retention-time, FWHM, and peak-base-width distributions, together with retention-time reproducibility and quantitative stability of internal standards |
| Mass accuracy | Precursor *m/z* and mass-deviation fields | Run-level and *m/z*-resolved mass-error assessment | Mass-error distributions, *m/z*-resolved median-error trends, and run-specific median references |
| Quantification quality | Quantity.Quality, Empirical.Quality, and PG.MaxLFQ.Quality | Joint visualization of DIA-NN-derived quantitative-quality fields | Run-resolved precursor-quality space and protein-group PG.MaxLFQ.Quality plots in relation to the number of distinct supporting precursors |
| Intensity and LFQ distribution | Precursor.Quantity and PG.MaxLFQ | Quantitative signal distributions across acquisitions | Precursor.Quantity density curves and protein-group PG.MaxLFQ boxplots for run-to-run comparison |
| Data completeness | Run identifiers, precursor identifiers, and quantities values | Observed run-precursor pairs relative to the inferred data space | Global and run-level tables of data-completeness percentages |
| Figures of Merit | Consolidated run-level QC metrics | Integration of identification, chromatography, mass, peptide, charge, and contaminant measures | Integrated table and CSV summarizing selected run-level results from the analytical modules |
| Ion mobility | Mobility, precursor m/z, and optional predicted mobility | Conditional density and trend assessment | Conditional run-level ion-mobility density plots and comparison of observed mobility with DIA-NN iIM trends when both fields are available |

### Reproducibility and Workflow Testing

Reproducibility was addressed by recording the scientific environment and preserving the computational context associated with report generation. For direct execution in R, renv.lock records the package dependencies required by the scientific workflow and supports reconstruction of the project environment. The portable Windows distribution provides the corresponding R runtime, package library, and Quarto installation, together with a graphical launcher generated from a PowerShell script using PS2EXE. The launcher collects the input paths selected by the user and initializes the bundled environment before passing execution to the same scientific runner used in the R-based mode. From this point onward, both modes share the same input-validation functions, analytical modules, and parameterized Quarto report, and no analytical calculation is implemented exclusively in the launcher. The generated HTML report preserves the date and time of execution together with the versions of QC-espresso, R, Quarto, and the relevant R packages, providing an execution-specific record of the computational environment.

The workflow was evaluated through automated tests maintained for software development and release verification. Focused tests assess individual analytical calculations, optional report sections, and multiprotein handling using controlled inputs and expected results. Integration and end-to-end tests examine the coordinated execution of the scientific modules and the generation of the complete report from example DIA-NN report.parquet and FASTA files. The resulting HTML reports and CSV exports were also inspected to confirm successful rendering and the presence of the expected outputs. The portable Windows distribution underwent additional execution testing to verify that the bundled environment and graphical launcher reproduced the expected report-generation workflow. Together, these procedures support the identification of regressions introduced by software modifications and are not required during routine use of QC-espresso. SHA-256 checksums are provided with the portable release archives as an additional integrity measure after download.

### Large Language Model Assistance during Software Development

Microsoft 365 Copilot assisted the development of QC-espresso through code standardization, debugging, and the development of selected testing and reproducibility procedures. The authors reviewed and, when appropriate, modified all suggestions before incorporating them into the software. Responsibility for the analytical design, validation criteria, acceptance of code changes, and final source code remained with the authors throughout development. Copilot was not used as an independent validation system. The source code and associated reproducibility resources are available in the project repository, allowing the final implementation to be inspected independently.

### Application to Representative Analytical Scenarios

To examine the practical application of QC-espresso, three DIA proteomics datasets processed with DIA-NN version 2.5.1 were selected to represent distinct analytical scenarios encountered in routine proteomics workflows. The examples address variation in sample composition and contaminant contribution, changes in chromatographic performance, and the relationship between mass accuracy and quantitative reliability. Each dataset was evaluated using its corresponding report.parquet file and FASTA database, with related measurements interpreted together and, where available, considered alongside the experimental records. A controlled three-proteome experiment with predefined sample compositions is presented in the Supporting Information, providing a complementary reference for the behavior of QC-espresso under analytically consistent conditions.

## Results and Discussion

Bottom-up proteomics workflows depend on consistent sample preparation and LC-MS/MS acquisition, as variation introduced at either stage can affect identification performance and quantitative accuracy. Sample preparation procedures, including protein extraction, enzymatic digestion, and sample clean-up, must be performed under conditions compatible with mass spectrometry analysis ^6^, while chromatographic and mass-spectrometric performance must remain sufficiently stable across acquisitions. Systematic examination of quality-control measurements can reveal changes associated with these different stages and guide further investigation before they affect downstream analyses ^7^.

### Sample Quality, Digestion Efficiency, and Contaminant Contribution

Sample preparation is a major source of variability in bottom-up proteomics because differences in individual preparation steps can affect subsequent stages and ultimately influence data quality. In particular, tryptic digestion is sensitive to the protein-to-enzyme ratio, reaction conditions, sample complexity, and enzyme quality, which can lead to substantial variation in the resulting peptide population ^6^.

Identification depth provides a first view of analytical recovery but cannot by itself explain whether an atypical result reflects digestion efficiency, sample composition, contamination, or expected biological variation. QC-espresso provides identification counts along with related sample-quality measurements, helping users recognize atypical samples and determine which aspects of the preparation workflow warrant further investigation. Evaluation across identification and quantitative levels is particularly important because sample-processing differences can propagate through the proteomics workflow with distinct effects on the final data quality ^8^.

Clear run-to-run variation was observed across the four panels, but the measurements did not always change in the same direction. Run 04 provides the clearest example: despite showing the highest missed-cleavage percentage, this sample also yielded the largest number of identified protein groups. Its quantitative profile was comparatively less dominated by individual proteins, and its contaminant contribution was among the lowest in the dataset. Thus, in this case, the elevated missed-cleavage rate did not coincide with reduced proteome recovery or with a generalized deterioration in sample composition. A different pattern was observed in runs 13 to 15, where lower identification depth was accompanied by a stronger predominance of selected signal components and markedly higher contaminant contributions.

Taken together, the panels reveal that sample quality is multidimensional, and that individual measurements may vary independently or converge on a common analytical pattern. Identification depth provides an overall view of proteome recovery, whereas missed cleavages offer a more specific view of digestion behavior. Signal composition indicates whether the recovered proteome remains broadly distributed or becomes dominated by a restricted set of proteins, and the contaminant profile helps assess whether this shift may be associated with exogenous or preparation-related material ^9^. The value of the combined visualization therefore lies in distinguishing isolated variation in one processing step from broader changes affecting both identification depth and sample composition.

## Chromatographic Performance across the Acquisition Series

The assessment of instrumental performance provides valuable information regarding the stability and reproducibility of LC-MS analyses. Variability in chromatographic systems may arise from column deterioration, changes in mobile-phase composition, flow instabilities, or instrument contamination, all of which can negatively affect acquisition quality ^7^. Such fluctuations are particularly problematic in large acquisition batches and longitudinal studies, where consistent performance across runs is essential. Furthermore, modern proteomics search algorithms rely heavily on retention time alignment to support confident peptide identification ^1^. Consequently, systematic shifts in chromatographic behavior may reduce identification performance and compromise quantitative consistency.

QC-espresso provides intuitive metrics for assessing chromatographic performance, a critical component of LC-MS quality control ^7^. These include the distribution of precursor identifications across the chromatographic gradient, which reflects sampling efficiency and separation performance, as well as peak-shape metrics such as full width at half maximum (FWHM) and peak-base-width, which provide measures of chromatographic resolution and peak performance.

For conventional trypsin-based bottom-up proteomics workflows, characteristic trypsin autolytic peptides ^10^ are monitored as internal reference markers of reproducibility and quantitative stability. Alternatively, endogenous internal standards ^11^ or synthetic calibration standards spiked into the samples may be supplied through the R scripts and incorporated into report generation, allowing QC-espresso to accommodate laboratory-specific practices and workflows. Internal peptide references are particularly important in DIA workflows, supporting RT normalization, assay library RT prediction for confident peptide identification, and accurate quantification through extracted ion chromatograms. QC-espresso automatically monitors the RT reproducibility and quantitative stability of these reference peptides, providing a convenient framework for continuous quality control of chromatographic performance and analytical consistency across acquisitions. The dataset illustrated in Figure 2 comprises multiple acquisition batches containing samples spiked with indexed retention time peptides (Biognosys iRT Kit). Following a failed acquisition (Run 40), the chromatographic column was replaced. Despite maintaining the same column specifications and conditioning the replacement column according to the manufacturer’s recommendations, QC-espresso report showed a clear reduction in identification performance following the exchange (Figure 2A). This reduction was accompanied by increased median FWHM values, indicative of peak broadening and reduced chromatographic resolution (Figure 2B). In addition, the RT of the reference peptides shifted by approximately five minutes after the column replacement (Figure 2C). RT subsequently returned to baseline values in the following acquisition batch (Run 51 onward). These observations demonstrate how chromatographic quality metrics establish useful references for retention time tolerances during data analysis and support consistent peptide identification and quantification across large-scale proteomics studies ^11^.

**Figure 1.**
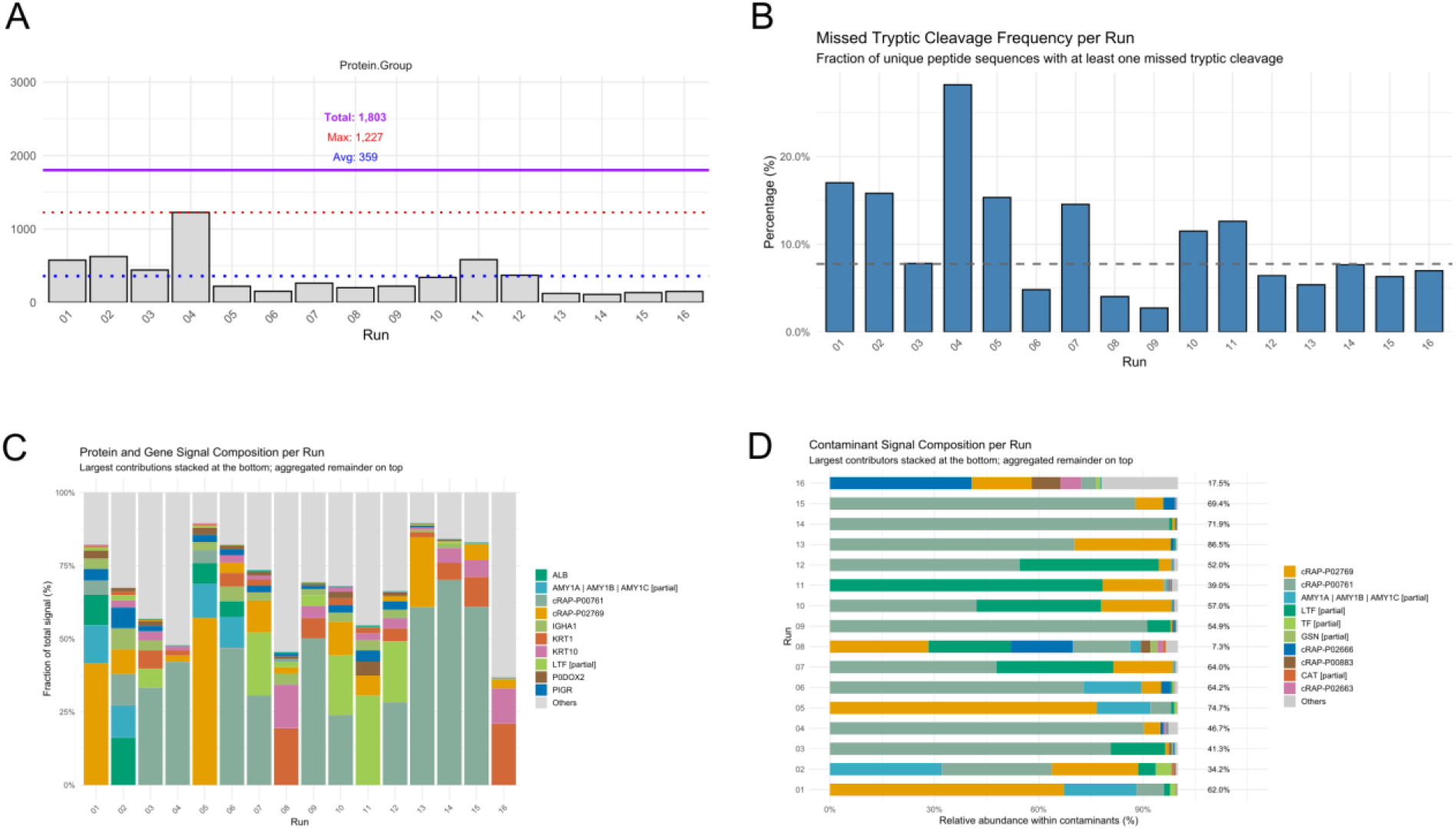
Sample quality, digestion efficiency, and contaminant contribution across runs. (A) Protein groups identified per run; lines indicate the mean, maximum per run, and total distinct groups across the dataset. (B) Percentage of unique peptide sequences with at least one missed tryptic cleavage; the dashed line indicates the median across runs. (C) Contribution of the 10 highest-signal protein or gene groups to the total quantitative signal; remaining groups are combined as “Others.” (D) Relative composition of contaminant signal based on recognized cRAP-or CON_ prefixes; values at right indicate the fraction of total quantitative signal assigned to contaminants in each run.

**Figure 2.**
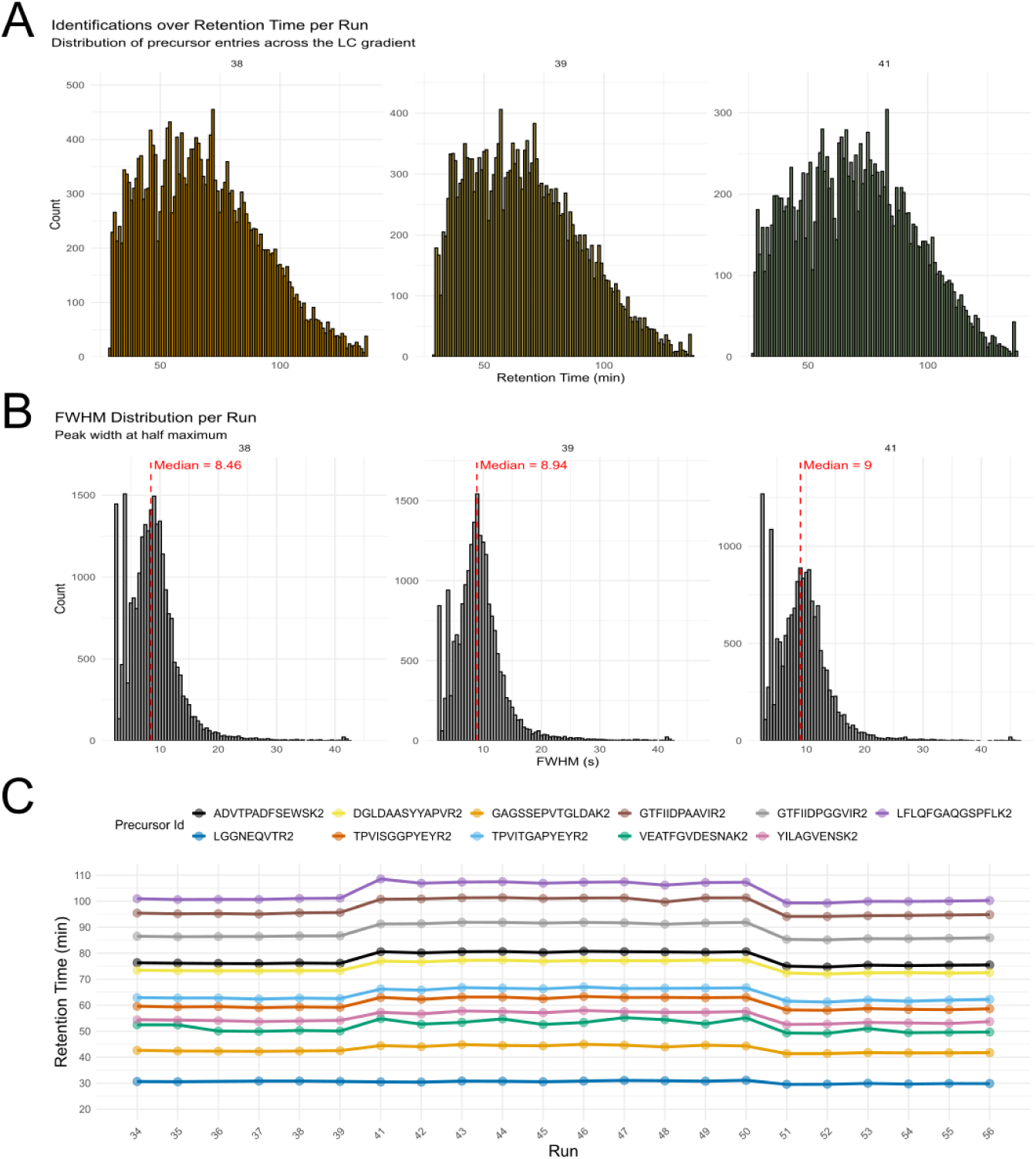
Chromatographic performance across acquisition batches. (A) Distribution of precursor identifications across the chromatographic gradient. (B) Full width at half maximum (FWHM) distribution of identified precursor features. (C) Retention time stability of internal standard peptides monitored throughout the acquisition sequence.

**Figure 3.**
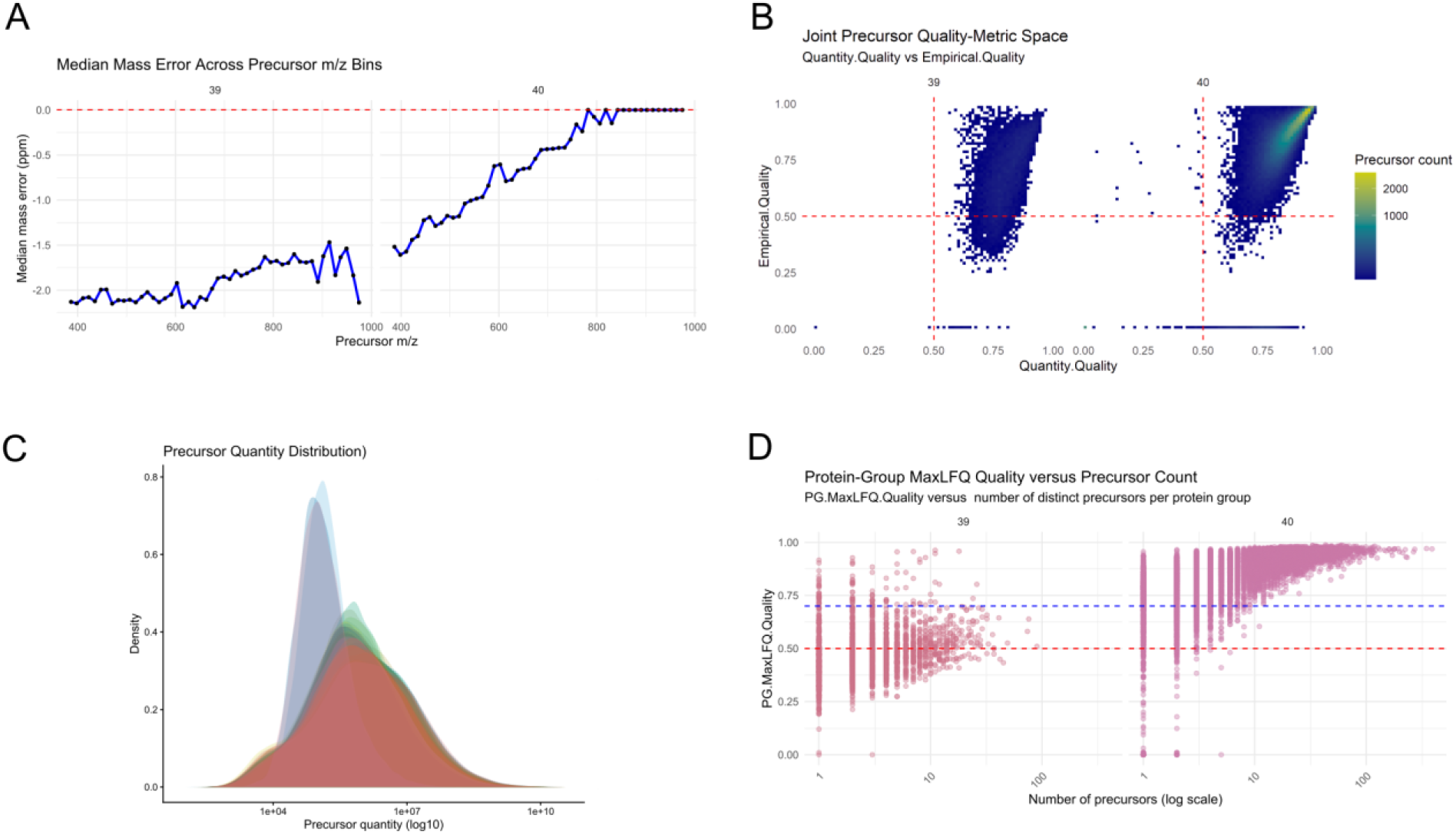
Acquisition performance and quantitative reliability across selected runs. (A) Median mass error across precursor m/z bins; the dashed line indicates 0 ppm. (B) Joint distribution of Quantity.Quality and Empirical.Quality, with color indicating precursor count; dashed lines at 0.5 serve as visual references. (C) Density distributions of positive Precursor.Quantity values on a logarithmic scale. (D) Protein-group PG.MaxLFQ.Quality in relation to the number of distinct supporting precursors; dashed lines at 0.50 and 0.70 serve as visual references.

### Acquisition Performance and Quantitative Reliability

In LC-MS-based proteomics, stable acquisition performance is essential for reliable identification and quantification. System-level metrics can reveal technical variation that may not be apparent from identification counts alone ^7^, with mass accuracy providing a particularly informative measure since systematic deviations can affect precursor matching and quantitative feature analysis ^12^. QC-espresso presents dedicated plots of DIA-NN-derived quantitative-quality measurements, allowing users to examine whether variation in precursor mass measurement coincides with broader changes in quantitative performance.

A consistent contrast was observed between runs 39 and 40 across the acquisition- and quantification-related measurements. In panel A, run 39 retained a more negative median mass error across the precursor m/z range, whereas the profile of run 40 progressively approached the 0 ppm reference at higher m/z values. The joint quantitative-quality distributions also differed between runs, with run 39 extending further toward lower Empirical.Quality values and run 40 showing a greater concentration of precursors in the higher Quantity.Quality and Empirical.Quality region (panel B). At the protein-group level, run 39 displayed a broader PG.MaxLFQ.Quality distribution, including numerous protein groups below the reference lines, whereas run 40 was predominantly concentrated at higher values, particularly for protein groups supported by multiple precursors (panel D). The overlapping precursor-quantity distributions in panel C provide an additional view of quantitative variation across the runs included in the analysis.

The concordance among the panels indicates that the contrast between runs 39 and 40 was not restricted to precursor mass measurement. The mass-error profile of run 39 coincided with a broader extension toward lower precursor- and protein-group quality values, while run 40 showed a mass-error profile approaching 0 ppm and a greater concentration of observations at higher quality values. The DIA-NN-derived QuantUMS-related fields provide uncertainty-related information that supports the assessment of quantitative reliability ^13^. Together, these measurements indicate when differences in acquisition performance extend across complementary levels of quantitative evidence, and the corresponding acquisition records and calibration status may warrant review. Collectively, the three case studies illustrate how a common QC-espresso report can accommodate analytical variation arising from sample quality and LC-MS/MS acquisition. By organizing DIA-NN-derived and sequence-based measurements within the same reporting structure, the workflow provides a consistent basis for examining the observed patterns in relation to their experimental context.

## Conclusions

QC-espresso transforms the extensive information available in DIA-NN outputs into an extended and scientifically inspectable record of experiment quality. Its contribution lies in making structured quality assessment accessible for routine use while retaining the analytical evidence, modular implementation, and computational context required for technical review. By bridging accessibility and scientific transparency, QC-espresso provides a practical foundation for standardized data quality documentation and the continued development of DIA proteomics workflows.

## Software and Data Availability

The QC-espresso source code, R modules, parameterized Quarto report, automated tests, example data, and renv.lock file are available at https://github.com/lanfredig/QC-espresso_DIA. The public example data comprise a DIA-NN report.parquet file of HeLa Pierce digest acquire in technical duplicates and its corresponding FASTA database, which can be used to execute and inspect the complete report-generation workflow. Version 1.3.1 was used for the analyses described in this study and is available through the repository Releases page, together with the portable Windows distribution. QC-espresso is distributed under the MIT License.

The case studies were derived from unpublished quality-control datasets generated at the mass spectrometry facility of the Brazilian Biosciences National Laboratory. These datasets were used exclusively to illustrate analytical quality-control scenarios and were not used to draw biological conclusions. Derived QC-espresso results for the cases presented in this study, including contaminant, chromatographic, quantitative-quality, and controlled benchmark examples, are available through the project repository.

## Author Contributions

G.P.L. and J.G.M. conceived and designed the study. G.P.L. led development and implementation of the tool. G.P.L. and J.G.M. conducted data analysis and data curation. All authors reviewed, revised, and approved the final manuscript.

## Acknowledgments

The authors thank André Zelanis and Mauricio Frota Camacho for testing QC-espresso and providing valuable feedback on its visual presentation and filtering alerts. The authors also thank Nilson Antonio da Rocha Coimbra for his advice on improving the robustness and organization of the software repository and Reynaldo Magalhães Melo for preparing the three-proteome LFQ benchmark samples used as a reference dataset in this study. The authors are grateful to Adriana Franco Paes Leme for supporting the development of this work and for providing the resources that enabled its completion. This study was conducted at the Brazilian Biosciences National Laboratory (LNBio), part of the Brazilian Center for Research in Energy and Materials (CNPEM).

## Abbreviations

DIA: data-independent acquisition
CSV: comma-separated values
FWHM: full width at half maximum
HTML: HyperText Markup Language
RT: retention time
LC-MS/MS: liquid chromatography–tandem mass spectrometry
LFQ: label-free quantification
QC: quality control

## Supporting Information

## Reference dataset

As a reference dataset, we employed the LFQ benchmark described by Navarro et al.^1^, consisting of a quality control sample composed of a three-species proteome mixture containing human HEK293, Saccharomyces cerevisiae, and Escherichia coli K12 protein lysates at defined proportions. The experimental design generated expected log₂ fold changes (B/A) of 0, -1, and +2 for the human, yeast, and bacterial proteomes, respectively (Figure 1). Samples were analyzed in technical triplicates to evaluate the reproducibility of both identification and quantification workflows.

**Figure S1.**
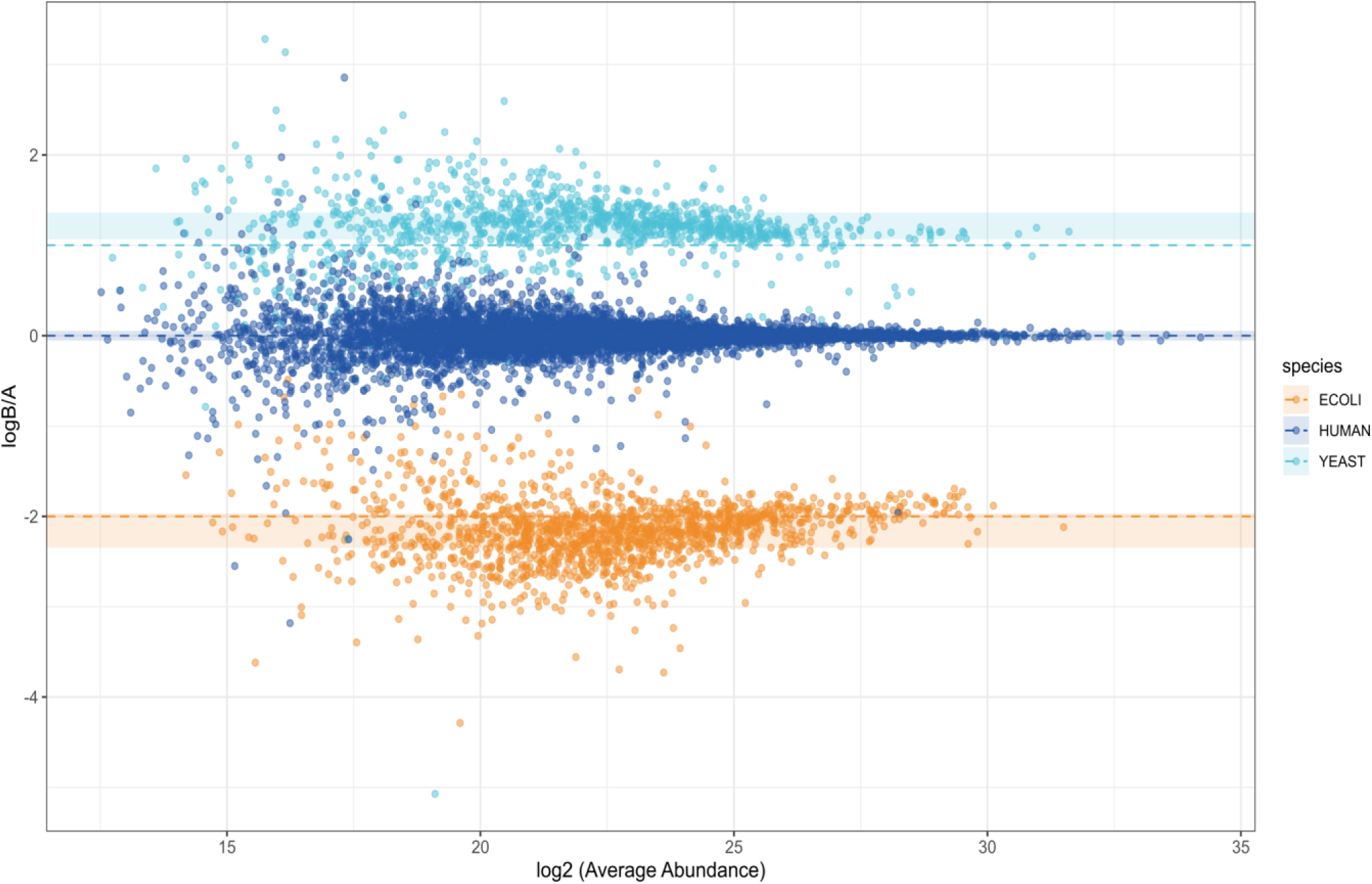
Recovery of expected protein abundance ratios in a three-species LFQ benchmark dataset. Observed protein log₂ fold changes between conditions A and B are shown as a function of average protein abundance. Dashed lines indicate the theoretical abundance ratios expected for human HEK293, *Saccharomyces cerevisiae*, and *Escherichia coli* K-12 proteins. Shaded regions represent the interquartile range of observed fold changes within each species group. Samples were analyzed in technical triplicates, 200ng injections, in a Orbitrap Astral with 2 *m/z* isolation window in data independent acquisition and 23 minutes gradient in an EASY-Spray 75µm x15cm PepMap Neo column. The analysis was performed using DIA-NN protein-group quantifications from technical triplicate measurements.

As illustrated in Figures 3A-E, the dataset exhibited robust analytical performance, characterized by stable chromatographic behavior, high mass accuracy, and consistent precursor, peptide, and protein identifications across replicates. A high reproducibility of precursor, peptide, and protein identifications (A), together with efficient proteolytic digestion (B) and low contaminant abundance (C), are expected from a robust sample preparation protocol. In addition, stable chromatographic performance (D) and low mass errors (E) contribute to highly reproducible quantitative performance across runs (F), ultimately reflected by the quality score metric (G). These results demonstrate that reliable identification and quantification can be achieved despite substantial changes in sample composition.

**Figure S2.**
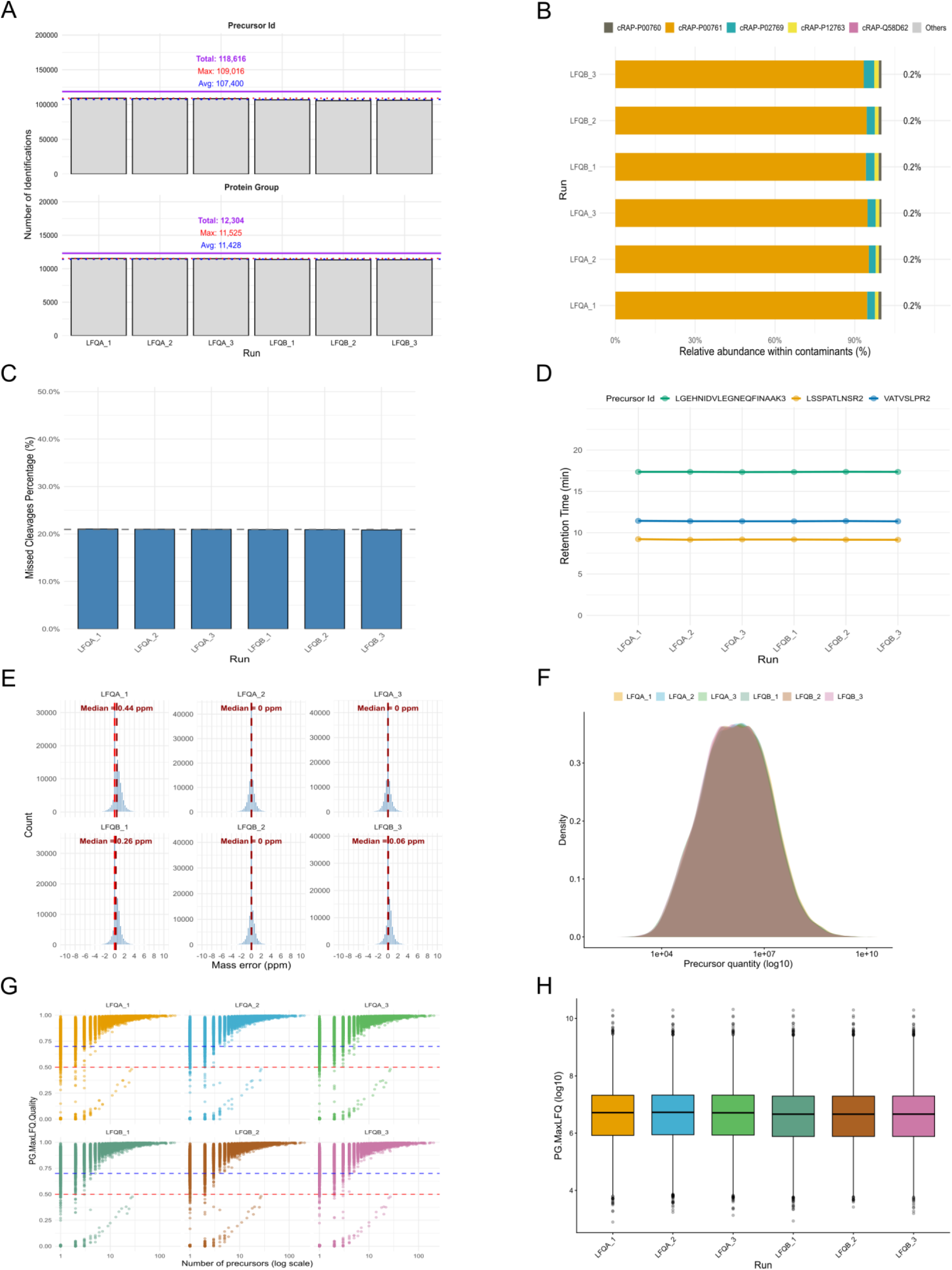
Application of QC Espresso to a three-species LFQ benchmark dataset. Representative quality control patterns expected from high-quality sample preparation and stable LC-MS instrumentation. (A) Reproducibility of peptide, and protein identification across runs. (B) Relative abundance of contaminant proteins. (C) Digestion efficiency metrics across runs. (D) Retention time distribution of characteristic tryptic autolytic peptides. (E) Mass error distribution across acquisitions. (F) Precursor-level quality metrics. (G) Protein-level quality metrics. (H) Density distribution of precursor MS2 quantity measurements. (I) Distribution of protein MaxLFQ intensities across runs.

